# A transcriptomic dataset for investigating the Arabidopsis Unfolded Protein Response under chronic, proteotoxic endoplasmic reticulum stress

**DOI:** 10.1101/2023.11.12.566746

**Authors:** Amélie Ducloy, Marianne Azzopardi, Caroline Ivsic, Gwendal Cueff, Delphine Sourdeval, Delphine Charif, Jean-Luc Cacas

## Abstract

The Unfolded Protein Response (UPR) is a retrograde, ER-to-nucleus, signalling pathway which is conserved across kingdoms. In plants, it contributes to development, reproduction, immunity and tolerance to abiotic stress. This RNA sequencing dataset was produced from 14-day-old *Arabidopsis thaliana* seedlings challenged by tunicamycin (Tm), an antibiotic inhibiting Asn-linked glycosylation in the endoplasmic reticulum (ER), causing an ER stress and eventually activating the UPR. Wild-type (WT) and a double mutant deficient for two main actors of the UPR (*INOSITOL-REQUIRING ENZYME 1A* and *INOSITOL-REQUIRING ENZYME 1B*) were used as genetic backgrounds in our experimental setup, allowing to distinguish among differentially-expressed genes (DEGs) which ones are dependent on or independent on IRE1s. Also, shoots and roots were harvested separately to determine organ-specific transcriptomic responses to Tm. Library and sequencing were performed using DNBseq™ technology by the Beijing Genomics Institute. Reads were mapped and quantified against the Arabidopsis genome. Differentially-expressed genes were identified using Rflomics upon filtering and normalization by the Trimmed Mean of M-value (TMM) method. While the genotype effect was weak under mock conditions (with a total of 182 DEGs in shoots and 195 DEGs in roots), the tunicamycin effect on each genotype was characterized by several hundred of DEGs in both shoots and roots. Among these genes, 872 and 563 genes were statistically up- and down-regulated in the shoot tissues of the double mutant when compared to those of WT, respectively. In roots of Tm-challenged seedlings, 425 and 439 genes were significantly up- and down-regulated in mutants with respect to WT. We believe that our dataset could be reused for investigating any biological questions linked to ER homeostasis and its role in plant physiology.

**SPECIFICATIONS TABLE:** 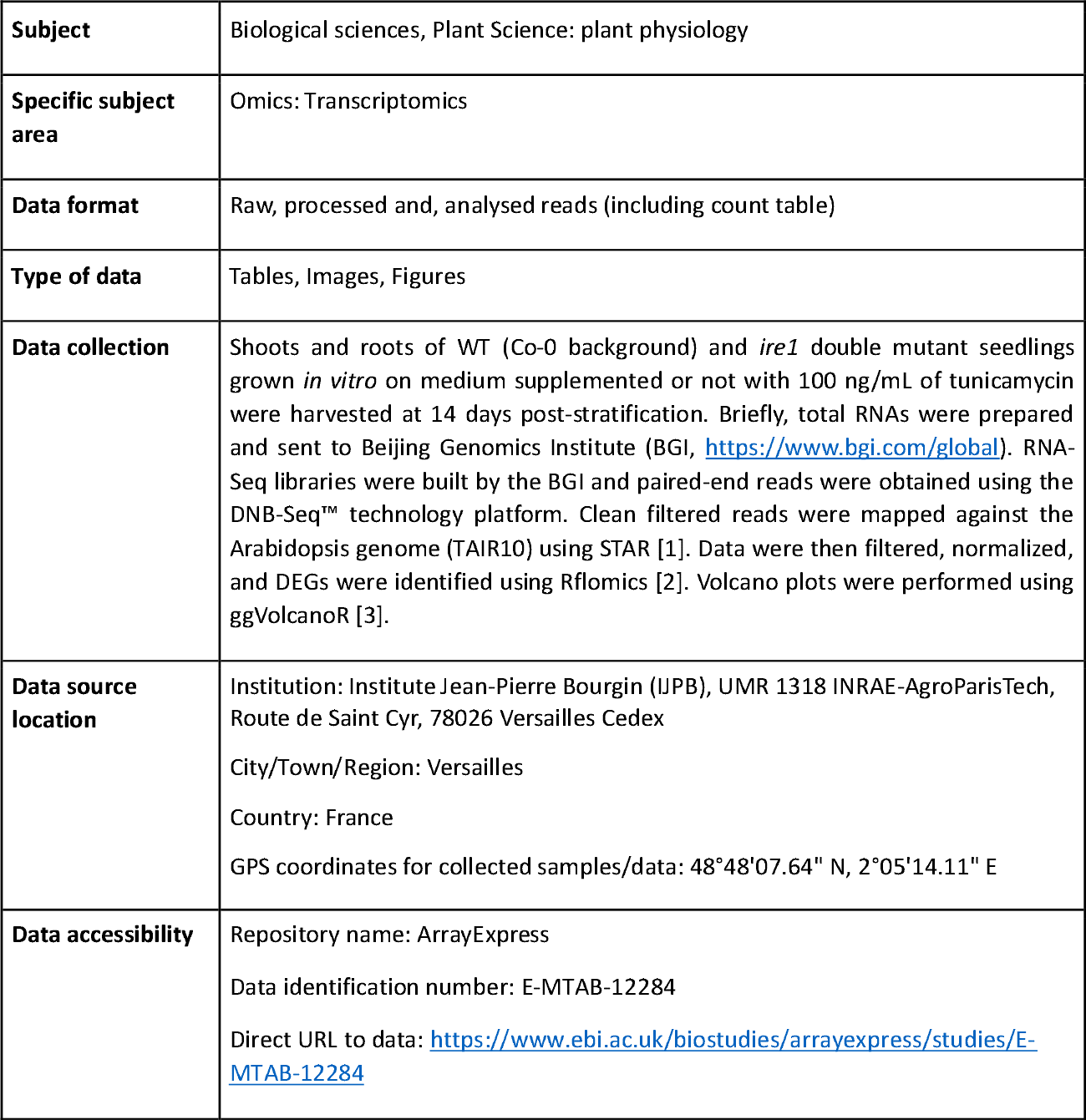

## VALUE OF THE DATA

These data contribute to our molecular understanding of the role of UPR under chronic ER stress, a situation that mimics environmental conditions where plants could be exposed to prolonged stresses, and trigger an acclimation response. This situation is poorly documented in the literature thus far. Our experimental design that consists of separately analyzing shoots and roots also allows for the identification of organ-specific transcriptional responses.

In addition to experts working on ER physiology and UPR, our transcriptomic dataset can benefit any researchers interested in plant response to stress, as ER stress is known to be triggered by numerous abiotic and biotic stresses. Since UPR is also invoked in root development and reproduction, our data can be reused by a wider part of the plant biologist community.

Our transcriptomic dataset can be used for identifying novel genes that play pivotal roles in ER stress response and tolerance, but also transcription factors that could be at the crosstalk between different stress signalling pathways in plants. These candidate genes could be further studied by reverse genetics to get a better understanding of their specific functions in regulatory networks.

## DATA DESCRIPTION

How the UPR differentially contributes at the organ level (shoots and roots) to plant tolerance under chronic ER stress needs to be documented. Here, we report on an organ-specific RNA sequencing dataset obtained from Arabidopsis WT (Col-0) and double *ire1* mutant (*ire1dm*) seedlings grown *in vitro* under chemically-induced ER stress condition. *ire1dm* provided by Pr. Koizumi (Osaka Prefecture University, Japan) were re-genotyped by PCR using left border and gene-specific primers (Fig. 1a-c). Based on PCR product sequencing (data not shown), T-DNA insertion in *IRE1B* gene was confirmed to locate upon the 1014^th^ nucleotide from the start codon (Fig. 1b), as expected from [4]. By contrast, insertion in *IRE1A* gene was relocated to position 2330 (counting from the start codon, Fig. 1a), which corresponds to SALK_002316 instead of SALK_018112. No or negligeable *IRE1A* and *IRE1B* transcripts corresponding to the genomic region downstream of the insertions could be detected by RT-qPCR in shoots and roots of *ire1dm*, under restful and stressful conditions (Sup. Fig. 1). To trigger a chronic proteo-toxic ER stress, tunicamycin was directly supplemented to the growth medium. Seeds were sown, stratified and germinated on supplemented medium, and seedling phenotype was determined at 14 days post-stratification (Fig. 1d). A Tm concentration of 100 ng/mL was selected as it has no impact on germination rates, irrespective of the tested genotypes (Fig. 1e). Moreover, this treatment provoked a 60% drop in fresh weight of both the shoots (Fig. 1f) and roots (Fig. 1g) of *ire1dm*. One such default in acclimation to Tm was not observed for WT and the re-genotyped, characrerized (Sup. Fig. 2) *bzip60-3* knock-out mutant (Fig. 1f,g), confirming previous studies [5] and validating our experimental conditions.

**Figure 1:**
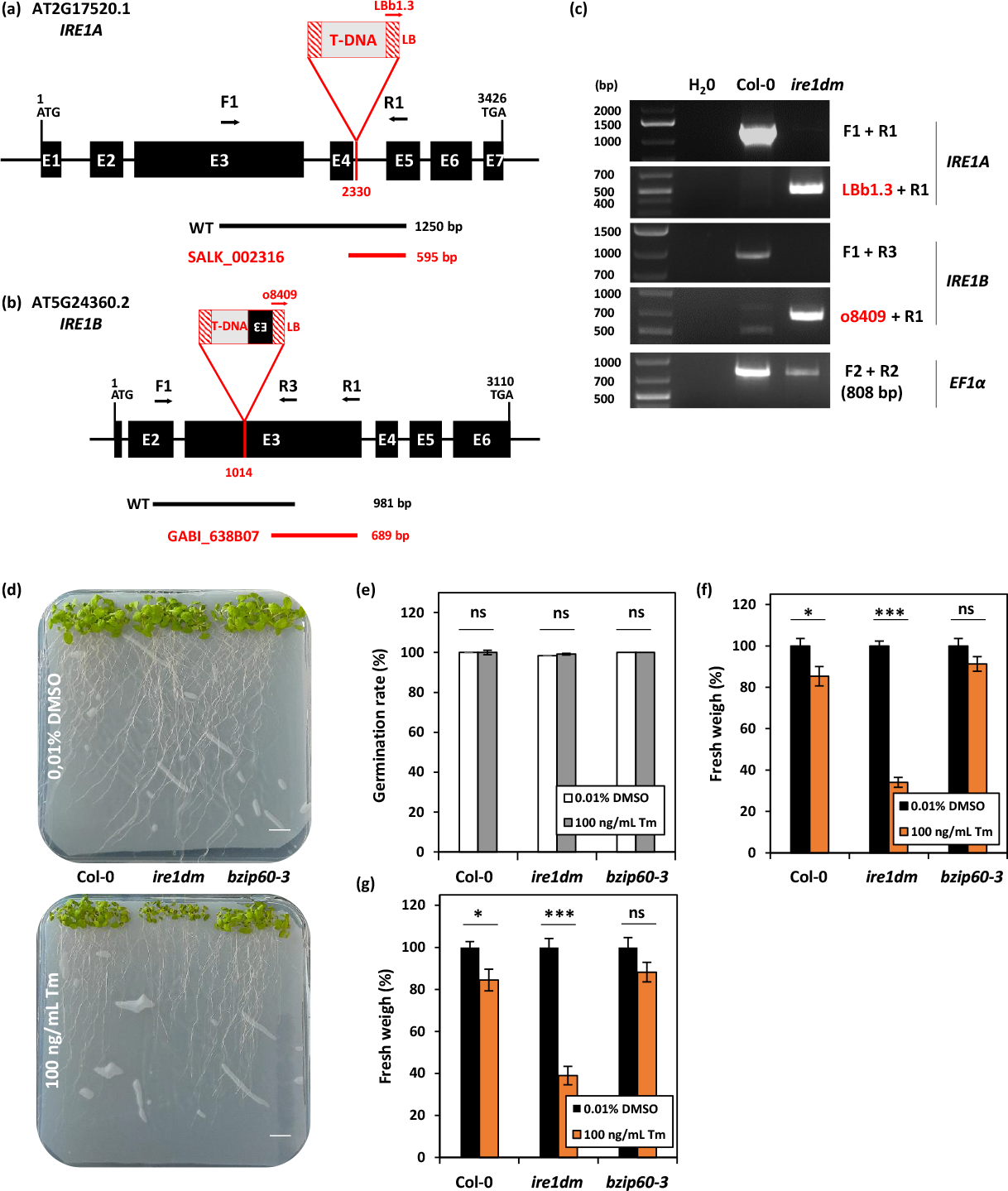
The double *ire1* mutant is hypersensitive to chronic ER stress induced by tunicamycin. Scheme of the T-DNA insertions in *IRE1A* (a) and *IRE1B* (b) genes of the double mutant (*ire1dm*). In these two panels, primer positions on the presented TAIR gene model, as well PCR product sizes, are indicated. PCR products shown were sequenced, allowing to relocate the exact position of the two T-DNA left border (LB) insertion sites (mentioned in red below the gene). Of note, part of the exon 3 of *IRE1B* was found to be inverted upstream of the T-DNA LB. (c) Representative results for the genotyping of the *ire1dm*. Combinations of primers used for PCR are indicated. (d) Representative photos of WT and mutant seedlings grown vertically for 14 days on tunicamycin-containing or DMSO-containing (mock) growth medium. Scale bar: 1 cm. (e) Germination rate of the three genotypes in mock and tunicamycin conditions. Mean ± SE for 8 to 32 biological replicates, each replicate including 15 seedlings. Student t-test was performed with α=0.05. ns, not significant. Average shoot (f) and root (g) fresh weight were quantified on 14-day-old seedlings for WT, *bzip60-3* and *ire1dm* genetic backgrounds. Mean ± SE for 10 biological replicates, each replicate including 15 seedlings. Student t-test was performed for assessing significant differences between mock and treatment. *, p-value<0.05; ***, p-value<0.001; ns, not significant.

RNA samples to be sequenced were prepared from shoots and roots of 14-day-old WT and *ire1dm* seedlings grown without (mock condition, named DMSO) or with Tm. Five biological replicates per modality were sequenced at the BGI using the DNB-Seq™ technology. On the whole, a total of 40 samples were sequenced and named after the genotype (WT or *ire1dm*), the condition (DMSO or Tm), the organ (shoot or root) and biological replicates (rep. number, see Table 1 for examples). All fastq raw data files, the count table and associated experimental information were deposited in the ArrayExpress repository under the data identification number E-MTAB-12284 (https://www.ebi.ac.uk/biostudies/arrayexpress/studies/E-MTAB-12284). Filtered reads were mapped to the *Arabidopsis thaliana* genome (TAIR10) using STAR [1]. Table 1 presents the quality of the mapping for all samples. Rflomics [2] was used to perform quality control (Fig. 2) and statistical analysis of gene expression for each organ, separately. Non- and low expressed genes (defined as those with an average count per million less than 1 in at least 4 samples) were filtered out and the TMM method was used to normalize gene expression (Fig. 2). In shoots, 14,172 out of 32,833 genes were discarded (18,661 remaining genes). In roots, 12,947 out of 32,833 genes were discarded (19,886 remaining genes). Remaining genes were considered as differentially-expressed when showing a Log_2_(FC)> 1 or a Log_2_(FC) <1 with an adjusted p-values < 0.05 (adjusted by Benjamini-Hochberg procedure). The number of identified DEGs in shoots and roots are shown in Volcano plots of Figure 3 and Figure 4, respectively.

**Table 1:** Rates of read mapping for all sequenced samples. Filtered reads were mapped against the TAIR10 version of Arabidopsis genome using STAR [1].

|  | Sample names | Total reads<br>(Millions) | Mapped reads<br>(Millions) | Mapped reads<br>(%) | Uniquely Mapped<br>(%) | Multi-Mapped<br>(%) | Unmapped<br>(too short)<br>(%) |
| --- | --- | --- | --- | --- | --- | --- | --- |
| SHOOTS | wt_dmso_shoot_rep2 | 33.71 | 33.57 | 99.60 | 98.50 | 1.00 | 0.40 |
|  | wt_dmso_shoot_rep4 | 33.64 | 33.47 | 99.50 | 98.40 | 1.10 | 0.48 |
|  | wt_dmso_shoot_rep5 | 33.65 | 33.53 | 99.60 | 98.60 | 1.10 | 0.37 |
|  | wt_dmso_shoot_rep6 | 33.64 | 33.47 | 99.50 | 98.50 | 1.00 | 0.53 |
|  | wt_dmso_shoot_rep7 | 33.53 | 33.36 | 99.50 | 98.30 | 1.20 | 0.50 |
|  | wt_tm_shoot_rep4 | 33.50 | 33.37 | 99.60 | 98.50 | 1.10 | 0.39 |
|  | wt_tm_shoot_rep5 | 33.66 | 33.50 | 99.50 | 98.40 | 1.10 | 0.49 |
|  | wt_tm_shoot_rep6 | 33.21 | 33.08 | 99.60 | 98.40 | 1.10 | 0.40 |
|  | wt_tm_shoot_rep7 | 32.85 | 32.75 | 99.70 | 98.30 | 1.40 | 0.30 |
|  | wt_tm_shoot_rep8 | 33.13 | 33.03 | 99.70 | 98.40 | 1.20 | 0.31 |
|  | ire1dm_dmso_shoot_rep3 | 28.07 | 27.98 | 99.70 | 98.50 | 1.20 | 0.32 |
|  | ire1dm_dmso_shoot_rep5 | 29.43 | 29.34 | 99.70 | 98.50 | 1.20 | 0.31 |
|  | ire1dm_dmso_shoot_rep6 | 32.53 | 32.42 | 99.70 | 98.40 | 1.30 | 0.33 |
|  | ire1dm_dmso_shoot_rep7 | 32.00 | 31.85 | 99.50 | 98.40 | 1.10 | 0.48 |
|  | ire1dm_dmso_shoot_rep8 | 32.96 | 32.81 | 99.50 | 98.40 | 1.10 | 0.47 |
|  | ire1dm_tm_shoot_rep1 | 33.35 | 33.23 | 99.60 | 98.00 | 1.70 | 0.35 |
|  | ire1dm_tm_shoot_rep2 | 33.56 | 33.45 | 99.70 | 98.40 | 1.30 | 0.31 |
|  | ire1dm_tm_shoot_rep4 | 33.39 | 33.25 | 99.60 | 98.40 | 1.20 | 0.40 |
|  | ire1dm_tm_shoot_rep6 | 33.44 | 33.32 | 99.60 | 98.30 | 1.30 | 0.36 |
|  | ire1dm_tm_shoot_rep7 | 31.95 | 31.80 | 99.50 | 98.20 | 1.30 | 0.46 |
| ROOTS | wt_dmso_root_rep1 | 32.84 | 32.58 | 99.20 | 97.80 | 1.40 | 0.80 |
|  | wt_dmso_root_rep2 | 33.39 | 33.14 | 99.30 | 97.90 | 1.40 | 0.72 |
|  | wt_dmso_root_rep3 | 27.96 | 27.78 | 99.40 | 98.00 | 1.30 | 0.65 |
|  | wt_dmso_root_rep4 | 32.93 | 32.67 | 99.20 | 98.00 | 1.20 | 0.80 |
|  | wt_dmso_root_rep7 | 33.54 | 33.33 | 99.40 | 98.10 | 1.30 | 0.61 |
|  | wt_tm_root_rep1 | 33.39 | 32.93 | 98.60 | 96.00 | 2.60 | 1.37 |
|  | wt_tm_root_rep2 | 33.07 | 32.85 | 99.30 | 98.00 | 1.30 | 0.67 |
|  | wt_tm_root_rep4 | 30.83 | 30.61 | 99.30 | 97.80 | 1.40 | 0.72 |
|  | wt_tm_root_rep6 | 33.04 | 32.76 | 99.20 | 97.80 | 1.30 | 0.86 |
|  | wt_tm_root_rep7 | 28.55 | 28.36 | 99.30 | 98.10 | 1.20 | 0.67 |
|  | ire1dm_dmso_root_rep1 | 33.12 | 32.90 | 99.30 | 97.80 | 1.50 | 0.66 |
|  | ire1dm_dmso_root_rep2 | 33.17 | 32.98 | 99.40 | 98.00 | 1.40 | 0.58 |
|  | ire1dm_dmso_root_rep3 | 33.13 | 32.95 | 99.50 | 98.10 | 1.40 | 0.54 |
|  | ire1dm_dmso_root_rep4 | 33.25 | 33.01 | 99.30 | 97.90 | 1.40 | 0.69 |
|  | ire1dm_dmso_root_rep8 | 33.02 | 32.78 | 99.30 | 97.90 | 1.40 | 0.71 |
|  | ire1dm_tm_root_rep1 | 31.78 | 31.58 | 99.40 | 98.10 | 1.30 | 0.62 |
|  | ire1dm_tm_root_rep2 | 33.20 | 32.95 | 99.20 | 97.90 | 1.40 | 0.73 |
|  | ire1dm_tm_root_rep3 | 33.13 | 32.96 | 99.50 | 98.20 | 1.40 | 0.48 |
|  | ire1dm_tm_root_rep4 | 33.16 | 32.94 | 99.30 | 97.90 | 1.40 | 0.66 |
|  | ire1dm_tm_root_rep6 | 28.99 | 28.84 | 99.50 | 97.60 | 1.90 | 0.51 |

**Figure 2:**
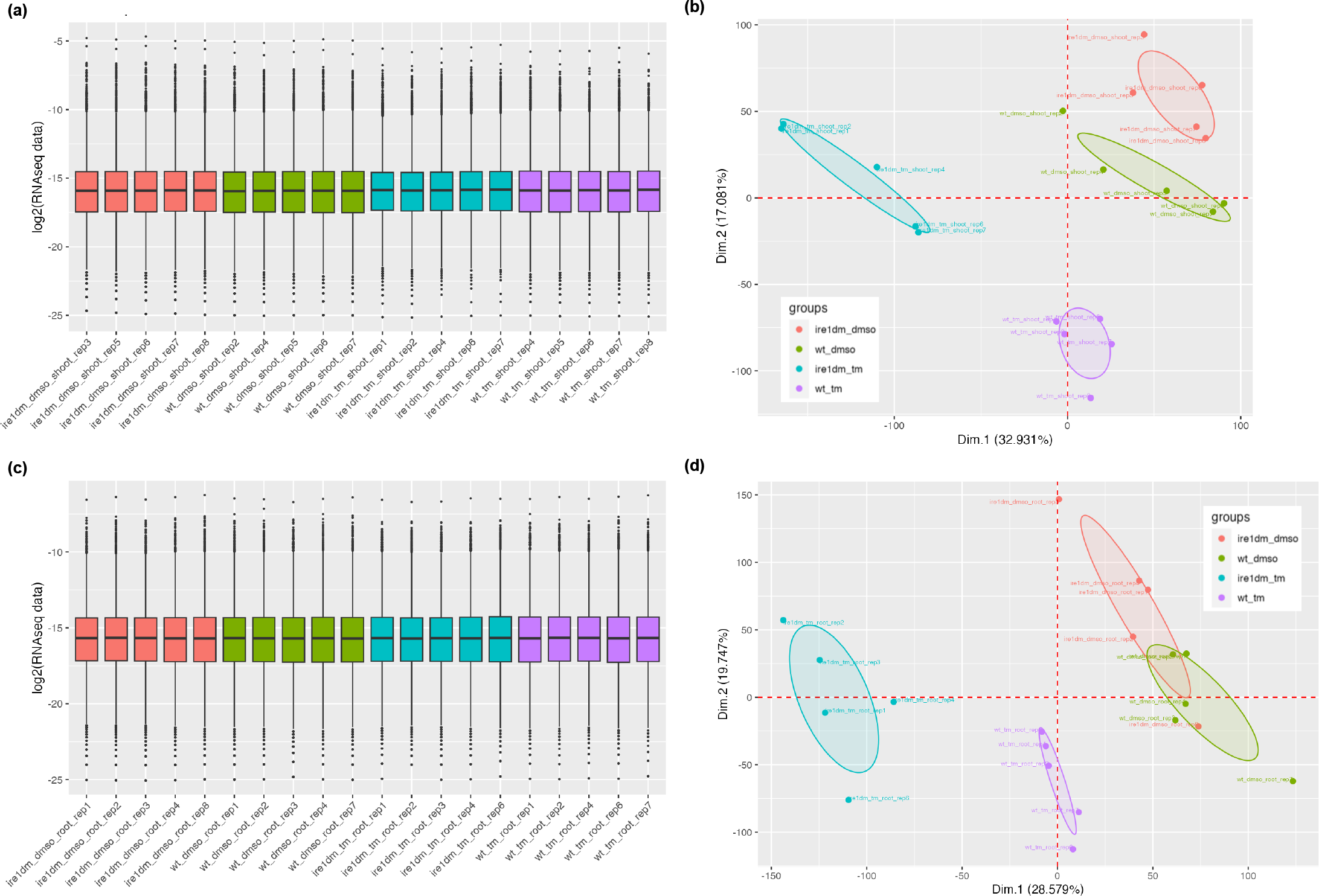
Quality control of the sequenced RNA biological replicates. The distribution of counts per sample upon filtering and normalization by the TMM method is presented for shoots (a) and roots (c). Repeatability of shoot (b) and root (d) samples was statistically analysed by Principal Component Analysis (PCA). Of note, the dimensions (Dim.) 1 and 2 explain 50% of the biological variability observed between modalities (genotype x treatment).

**Figure 3:**
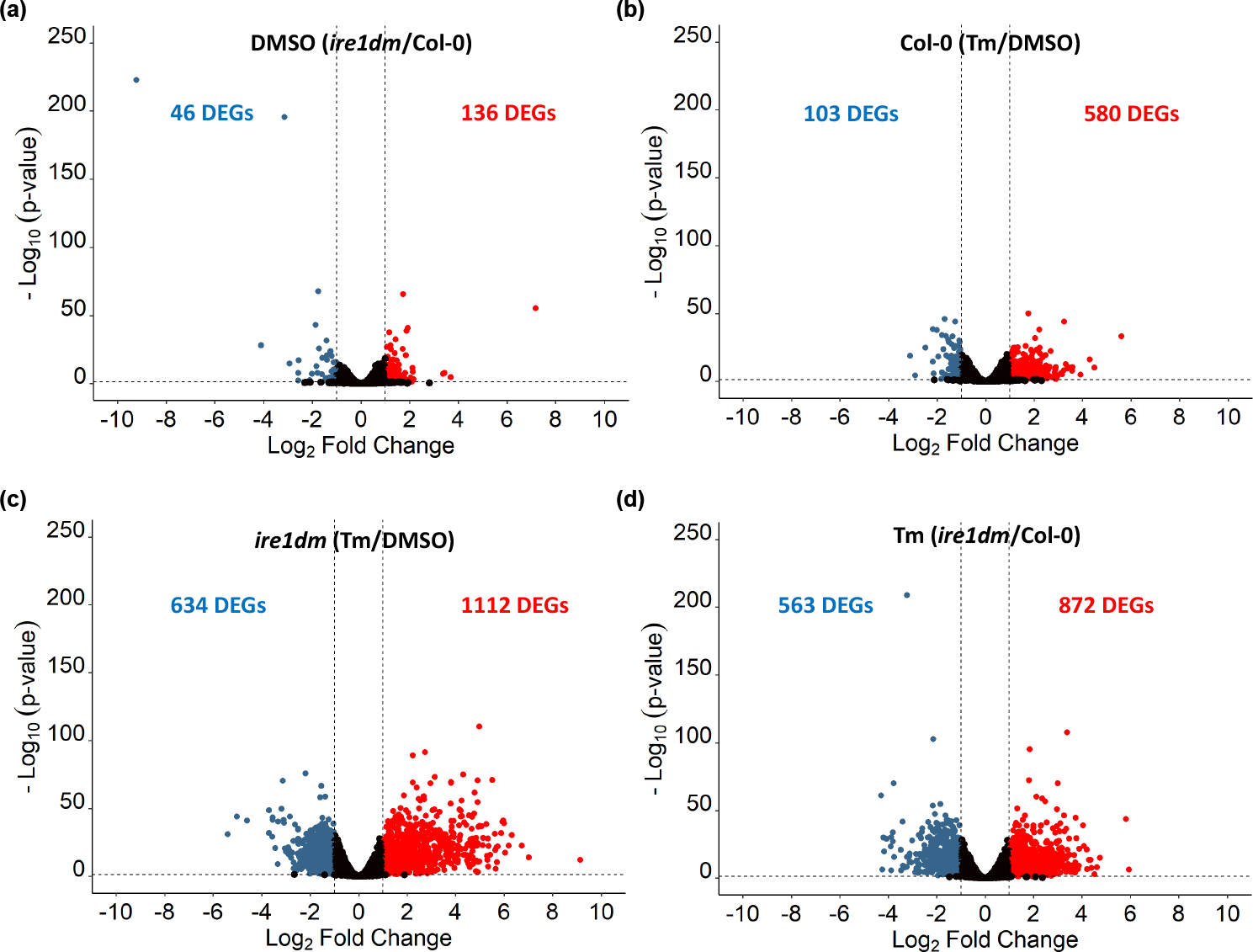
Differentially-expressed genes across conditions and genotypes in shoots. Volcano plots showing the genotype effect in mock (DMSO) conditions (a), the tunicamycin effect in WT (b), the tunicamycin effect in *ire1dm* (c), and the combination of the genotype and tunicamycin effects (d). The red and blue colors indicate up- and down-regulated genes, respectively. Significant DEGs are shown for a Log_2_(Fold Change) > 1 or <1 and an adjusted p-value < 0.05 (Benjamini-Hochberg procedure).

**Figure 4:**
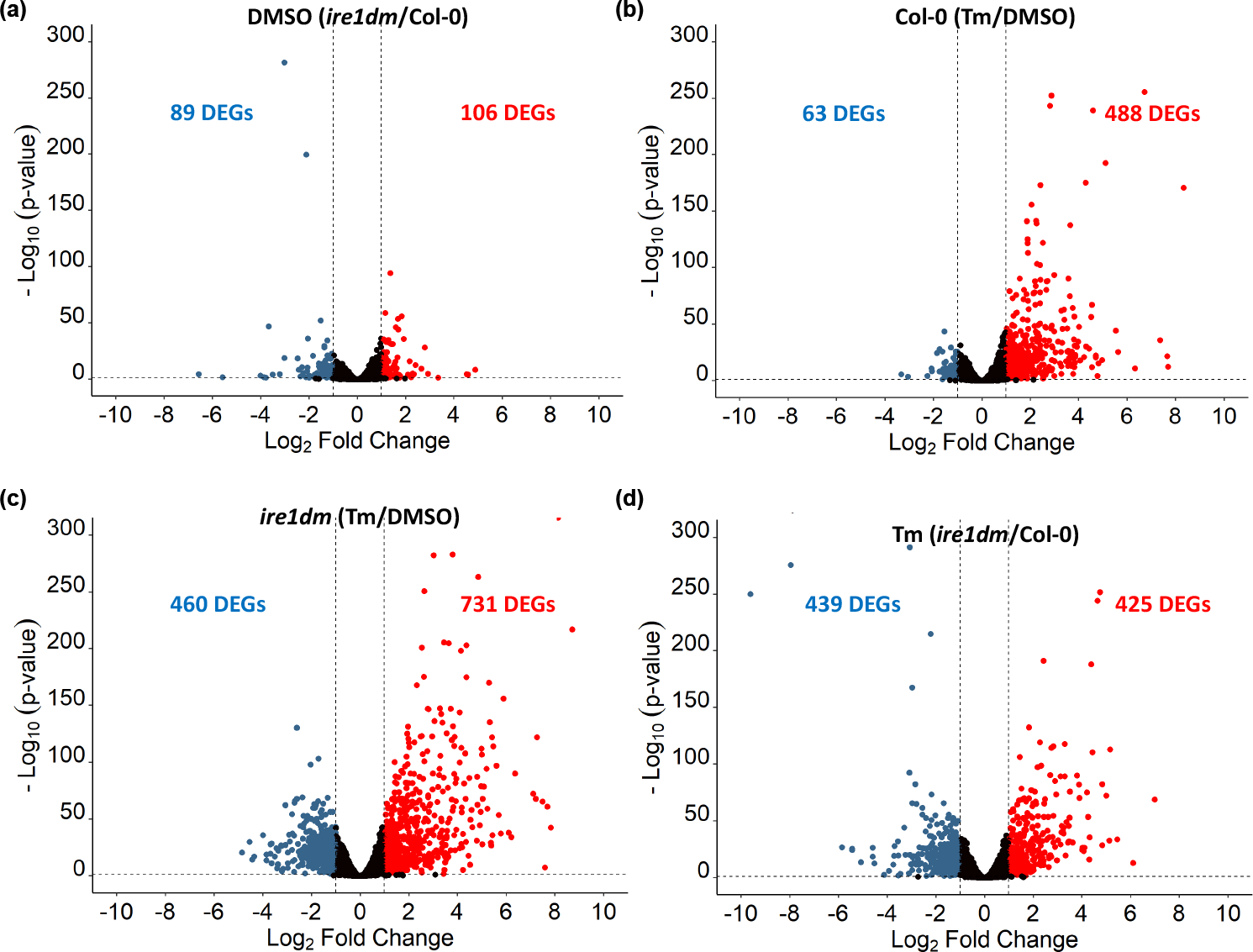
Differentially-expressed genes across conditions and genotypes in roots. Volcano plots showing the genotype effect in mock (DMSO) conditions (a), the tunicamycin effect in WT (b), the tunicamycin effect in *ire1dm* (c), and the combination of the genotype and tunicamycin effects (d). The red and blue colors indicate up- and down-regulated genes, respectively. Significant DEGs are shown for a Log_2_(Fold Change) > 1 or <1, and an adjusted p-value < 0.05 (Benjamini-Hochberg procedure).

## EXPERIMENTAL DESIGN, MATERIALS AND METHODS

### Plant material and growth conditions

Three genotypes were used in this work: the single mutant *bzip60-3*, the double mutant *ire1a ire1b* [4] and the corresponding WT genetic background, Columbia-0 (Col-0). Mutant alleles were GABI_326A12 (*bzip60-3*), SALK_002316 (*ire1a*) and GABI_638B07 (*ire1b*). Sterilized seeds were directly sown and germinated on nylon mesh laying on a modified, solid Murashige & Skoog medium (DU0742.0025, Duchefa, pH=5.8) supplemented with Ca(NO_3_)_2_ at a final concentration of 2 mM and 1.2% (w/v) Phytoblend (PTP01, Caisson Labs). Upon stratification (4°C in the dark for 3 days), seedlings were grown vertically in a growth chamber (Sanyo, MLR-350H model) for 14 days, in long day conditions (16h light at 22°C, 8h night at 21°C) with a light intensity of 150 μmoles of photons.m^-2^.s^-1^. For Tm treatment, the compound (T7765, Sigma-Aldrich) was directly added to the growth medium at a final concentration of 100 ng/mL. Since Tm was dissolved in dimethylsulfoxide (DMSO) as solvent, control plates were supplemented with a volume of DMSO equivalent to that of Tm (0.01%, v/v). Mutant and WT genotypes were always grown side by side in plates. Independent biological replicates (roughly composed of 15-20 seedlings per genotype) were always grown on different plates.

### Genotyping of mutants

The single *bzip60-3* and double *ire1* mutants were re-genotyped by PCR (Fig. 1, Sup. Fig. 2) and PCR products were sequenced to precisely relocate the insertions (data not shown). Genomic DNA used as PCR matrix was extracted from leaf material according to [6]. Mutant alleles were genotyped using left border specific primers (LBb1.3 for SALK and o8409 for GABI-Kat lines) and gene specific primers described in Supplemental Table 1.

### Phenotyping of mutants

For fresh weight quantification, shoots and roots of 14-day-old seedlings were separated using a scalpel and weighted using a precision balance (10 biological replicates per genotype, each including 15 seedlings). Shoot and root weights were normalized by the number of seedlings per biological replicate, and expressed in percent. The germination rate was measured at 5 days post-stratification considering seeds as germinated when cotyledons were visible (8-32 biological replicates, each including 15 seedlings), and it was expressed in percent.

### RNA extraction

Shoots and roots of 14-day-old seedlings were separated using a scalpel, directly frozen in liquid nitrogen and stored at -80°C until extraction. Starting from 80 to 100 mg fresh weight per sample, total RNAs were extracted using the RNeasy Plant Mini kit (#74904, QIAGEN) following manufacturer’s instructions. RNAs were solubilized in DNAse-free water, quantified with a SPECTROstar Nanodrop (BMG LABTECH) and stored at -80°C until sequencing or reverse transcription. Again, independent biological replicates were always grown on different plates.

### Quantification of *IRE1 and bZIP60* gene expression by RT-qPCR

To get rid of DNA contaminations, total RNAs (1μg/sample) were incubated with DNase I (RNase-free, EN0525, ThermoFisher Scientific) at 37°C for 30 min. DNase I was subsequently inactivated at 65°C for 10 min. RNAs were then reverse transcribed in the presence of RNaseOUT (#10777019, ThermoFisher Scientific) with SuperScript II reverse transcriptase using oligo(dT) as primers, following the manufacturer’s instructions (#18064014, ThermoFisher Scientific). Samples were treated with RNase H (EN0201, ThermoFisher Scientific), and single strand cDNAs subsequently stored at -20°C until further use.

For qPCR, pure cDNAs were diluted 50 times and 2.5 μL of the diluted solution were used as matrix (final reaction volume of 5 μL). Reaction mix were prepared using SYBR Green Supermix (#1708880, Biorad). The final concentration of each primer was 2.5 μM. qPCR was run using the QuantStudio 5 machine (A28140, Applied Biosystems) with the following program: initial denaturation at 95°C for 3 min, followed by 40 cycles of denaturation at 95°C for 15s and combined annealing/extension at 63°C for 50s. Data were analyzed with QuantStudio Design and Analysis software (version 2.6). Data normalization was performed with three reference genes (*ACT2, EF1α* and *RHIP1*), according to the Cqmin method previously described [7]. Four independent biological replicates, each including two technical replicates, were analyzed for each gene. Prior to gene expression quantification, primers efficiency was determined using cDNA serial dilution (starting from 1:10 solution, diluted seven times by 5). qPCR primer sequences and efficiencies are provided in Supplemental Table 2.

### RNA sequencing

Five micrograms of total RNAs were treated with DNase I (RNase-free, EN0525, ThermoFisher Scientific). DNase I was heat-inactivated and treated RNAs were precipitated overnight at 4°C using an equal volume of 4M LiCl solution. High molecular weight RNAs were pelleted by centrifugation at 8,000 x g for 30 min at 4°C. Pellets were resuspended in 50 μL of RNase-free water and 150 μL of cold ethanol was added. Samples were incubated for 1 h at -80°C, centrifuged and the pellet was rinsed once with 70% cold ethanol before drying. RNAs were resuspended in RNase-free water and their integrity was checked on agarose gel (data not shown). Prepared RNA samples were then sent to the Beijin Genomics Institute (BGI, Hong Kong, https://www.bgi.com), and their quality and concentration were checked using bioanalyzer of the BGI in-house facilities. Seven to eight RNA samples were prepared for each modality (condition x genotype). Based on the RIN value and ratio RNA 28S/RNA 18S, the best five samples for each modality were chosen for sequencing. Sequenced shoot RNA samples displayed RIN values ≥ 6.7 and a ratio RNA 28S/RNA 18S ≥ 1.5 (data not shown). Sequenced root RNA samples displayed RIN values ≥ 7.7 and a ratio RNA 28S/RNA 18S ≥ 1.7 (data not shown). BGI was in charge of the cDNA library building (Eukaryotic Strand-specific mRNA) and sequencing (paired-end reads of 150bp, 30M clean reads per sample) using the DNA NanoBall Technology (DNB-Seq™). After sequencing, raw reads were filtered by the BGI to reach a Phred+33 fastq quality score.

### Analysis of RNA-Seq data

Filtered reads were mapped to the *Arabidopsis thaliana* genome (TAIR10) using STAR [1], with default parameters. Gene expression was quantified using the htseq-counts program [8] taking into account that the libraries are strand-specific (--stranded=“reverse”), counting reads by exon (-- type=“exon”) and only considering those with a mapping quality up to 20 and specific to a gene annotation (--mode=“union”). Rflomics R package and shiny web application [2] were used to perform quality control and statistical analysis of gene expression for each organ (shoot and root) separately, with the same methods and parameters. Non- and low expressed genes (defined as those with an average count per million less than 1 in at least 4 samples) were filtered out and the edgeR::TMM method was used to normalize gene expression. Quality controls was performed by checking distribution of counts before and after these two steps and by checking that technical replicates groups together using Principal Component Analysis (ensuring that technical variability is low with regards to biological outcome). Differential expression analysis was carried out for each selected contrasts (DMSO (*ire1dm*/Col-0); Tm (*ire1dm*/Col-0); Col-0 (Tm/DMSO) and *ire1dm* (Tm/DMSO)) thanks to the edgeR package. Parameters of the generalized linear model was first estimated thanks to the edgeR::estimateGLM* functions and the model was then fit using the edgeR::glmFit function. Likelihood ratio tests (LRT) were then performed for each contrast thanks to the edgeR::glmLRT function. P-values were adjusted by Benjamini & Hochberg procedure. Genes were considered as differentially-expressed when showing a Log_2_(FC)> 1 or a Log_2_(FC)< 1, with an adjusted p-values < 0.05.

## LIMITATIONS

Not applicable.

## ETHICS STATEMENT

The authors have read and follow the ethical requirements for publication in Data in Brief and confirm that the current work does not involve human subjects, animal experiments, or any data collected from social media platforms.

## CRediT AUTHOR STATEMENT

**Amélie Ducloy**: Investigation, Validation. **Marianne Azzopard**i: Investigation, Validation, Visualization, Review & Editing. **Caroline Ivsic:** Investigation. **Gwendal Cueff**: Formal analysis, Data curation. **Delphine Sourdeval:** Investigation. **Delphine Charif:** Data curation, Review & Editing. **Jean-Luc Cacas**: Conceptualization, Investigation, Visualization, Supervision, Writing-Original Draft/Review & Editing, Funding Acquisition.

## ACKNOWLEDGEMENTS

We wish to warmly thank Pr. Nozomu Koizumi (Osaka Prefecture University, Japan) for sharing his genetic resources, including the double *ire1* mutant. We also warmly thank Pr. Stephen Howell (Iowa State University, USA) for providing the *bzip60-3* mutant. We thank Dr. Ivan Le Masson (UMR Agronomie, INRAe, France) for protocol and tips about RNA precipitation by lithium chloride. We thank Pr. Loïc Rajjou (AgroParisTech, University Paris-Saclay, France) for stimulating discussions. We also would like to thank Pr. Marie-Noëlle Bellon-Fontaine and Dr. Aurélie Baliarda (AgroParisTech, University Paris-Saclay, France) for their kind support. RNA sequencing was supported by AgroParisTech [“Fédérateur” funding call, grant AOSV_001_4PBA].

## Supplemental figure legends

**Supplemental figure 1: Organ-specific expression of** *IRE1* **genes in WT and double mutant seedlings challenged by tunicamycin**. *IRE1A* (a) and *IRE1B* (b) gene expression was quantified by qPCR as described in the section “Experimental Design, Material & Methods” using primers reported in Supplemental Table 2. Primers used for that purpose were matching sequences corresponding to the genomic region localized downstream of the insertions. Residual expression (%) for both genes *in ire1dm* when compared to mock WT are indicated in bubbles. Statistical differences (n=4 biological replicates) were assessed by one way ANOVA and a post-hoc Tuckey HSD test with α=0.05. ns, not significant. nd, not detected (no amplification and/or Cq>35).

**Supplemental figure 2: Molecular characterization of the** *bzip60-3* **mutant**. (a) Scheme of the T-DNA insertion in *bZIP60* gene in the mutant. Primer positions on the presented TAIR gene model, as well PCR product sizes, are indicated. PCR products shown were sequenced, allowing to relocate the exact position of the T-DNA left border (LB) insertion sites (mentioned in red below the gene). (b) Representative results for the genotyping of the *bzip60-3* mutant. Combinations of primers used for PCR are indicated. (c, d) Organ-specific expression of *bZIP60* gene in WT and mutant seedlings challenged by tunicamycin. *bZIP60* unspliced (a) and spliced (b) transcripts (noted *bZIP60u* and *bZIP60s*, respectively) were quantified by qPCR as described in the section “Experimental Design, Material & Methods” using primers reported in Sup. Table 2. Primers used for that purpose were matching sequences corresponding to the genomic region localized downstream of the insertion. Statistical differences (n=4 biological replicates) were assessed by one way ANOVA and a post-hoc Tuckey HSD test. * significant with α<0.05, ** significant with α <0.01, ns, not significant. nd, not detected (no amplification and/or Cq>35).

**Supplemental table 1:**
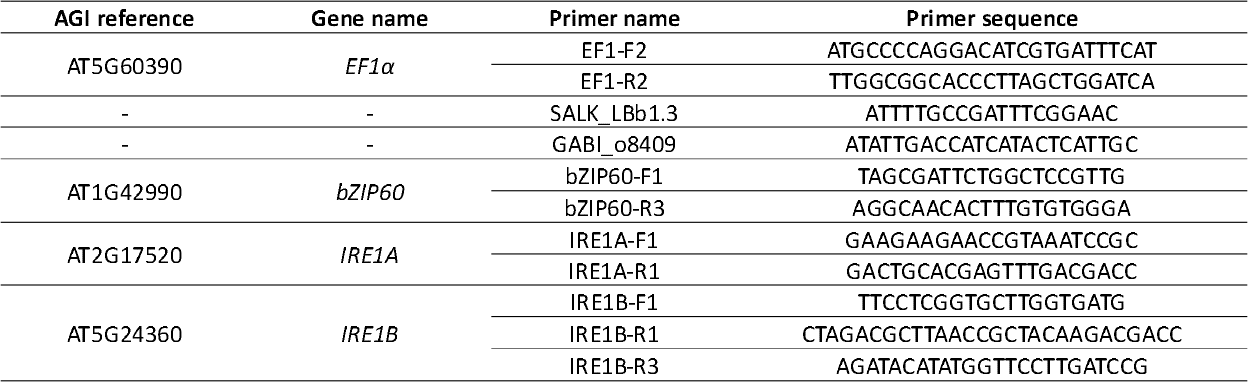
Left border, house-keeping gene- and gene-specific primers used for genotyping *bzip60-3* and the double *ire1* mutants. AGI: Arabidopsis Genome Initiative.

**Supplemental table 2:**
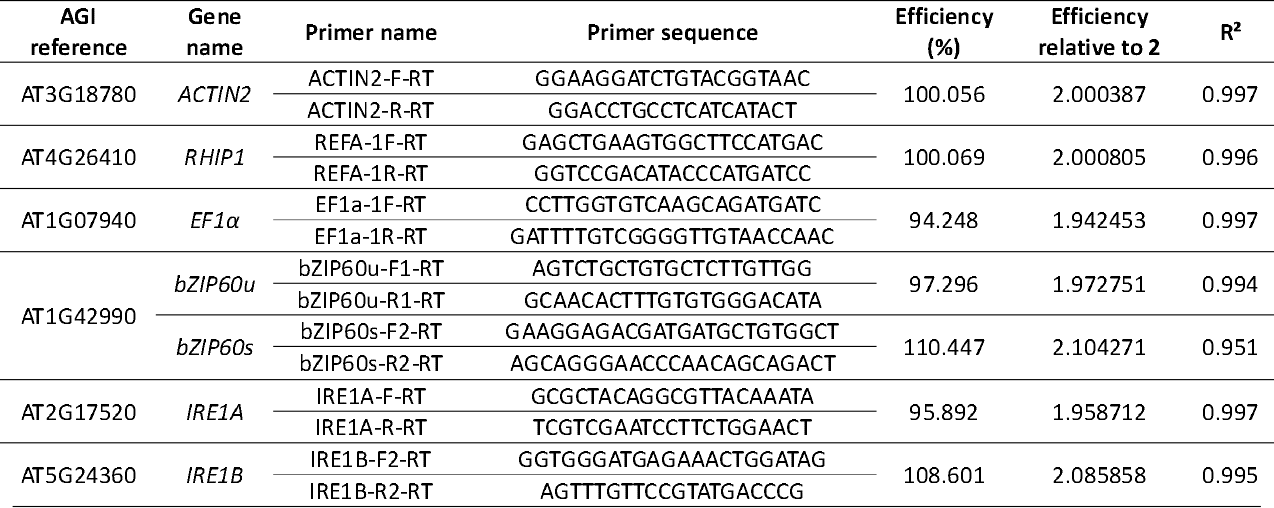
Primers used for quantifying the expression of *bZIP60* and *IRE1* genes by RT-qPCR. For quantifying *bZIP60* expression, two pairs of primers were used: one allowing to amplify cDNAs corresponding to the unspliced mRNAs (noted bZIP60u) and another one allowing to amplify cDNAs corresponding to the spliced mRNAs (noted bZIP60s). The correlation coefficient R^2^ obtained when determining primer efficiency is provided. AGI: Arabidopsis Genome Initiative.

