## Supplementary figures and images for "A transcriptomic dataset for investigating the Arabidopsis Unfolded Protein Response under chronic, proteotoxic endoplasmic reticulum stress"

### Supplemental figures 1 & 2

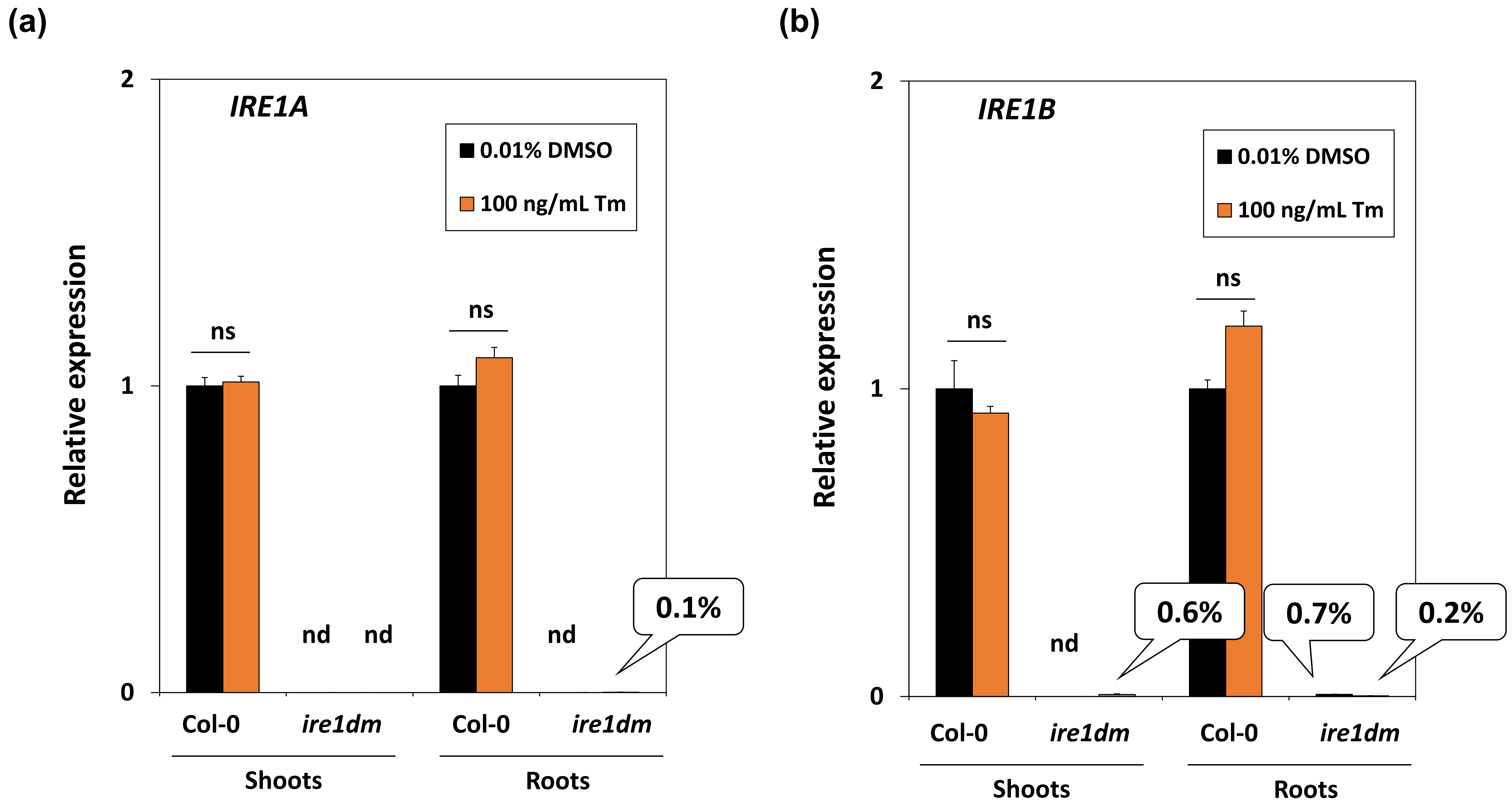


**Supplemental figure 1**

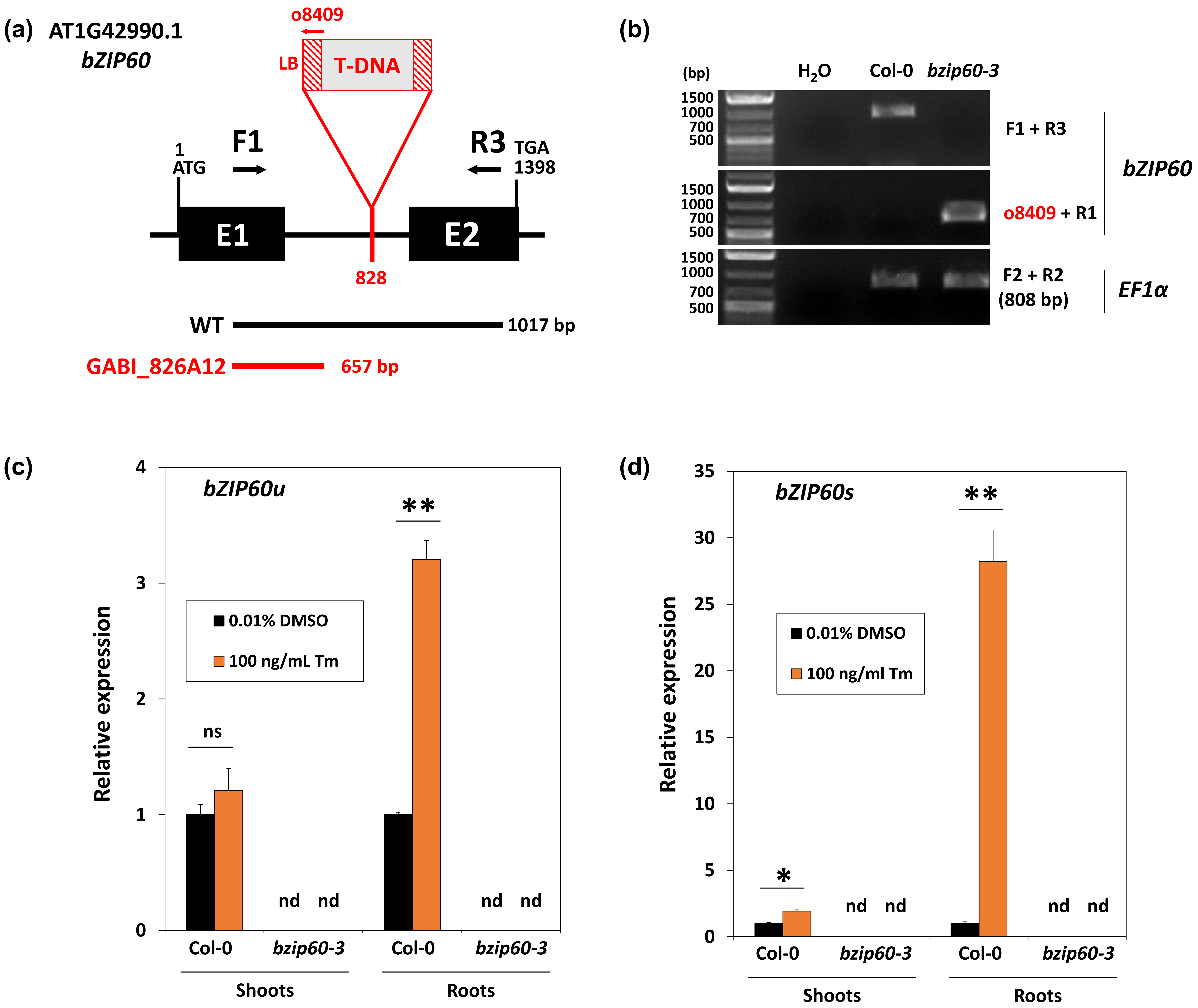


**Supplemental figure 2**
